# Early life stress modulates behavioral sensitivity to alcohol and promotes escalation of alcohol drinking

**DOI:** 10.1101/2025.06.27.661986

**Authors:** Allison R. Morningstar, Olivia S. R. Ledbury, Angeline C. Yu, Haniyyah Sardar, Ethan T. Rogers, Isaac F. Kandil, R. Nicolas Fajardo, Max E. Benabou, William J. Giardino

## Abstract

Adverse childhood experiences significantly increase the risk of developing alcohol use disorder (AUD) in adulthood. We used a model of combined limited bedding/nesting and maternal separation (LBN+MS) in C57BL/6J background mice to investigate how early life stress (ELS) modulates behavioral sensitivity to alcohol, long-term alcohol drinking patterns, and the effects of alcohol on social behaviors. Our findings reveal that ELS increased sensitivity to the stimulatory locomotor effects of alcohol (1.75 g/kg) selectively in females and reduced sensitivity to the sedative effects of alcohol (4.0 g/kg) particularly in males. This pattern of enhanced stimulation and diminished sedation is consistent with phenotypes observed in human subjects at high risk for developing AUD. ELS also significantly enhanced escalation of voluntary alcohol intake and preference over eight weeks of two-bottle choice intermittent access drinking particularly in males. Additionally, social behavior assessments revealed that ELS impaired sociability selectively in females with a history of alcohol drinking, highlighting the detrimental interactive effects of ELS and alcohol exposure on adaptive behaviors. These results underscore the complex interplay between ELS, alcohol responses, and sex differences, suggesting that ELS creates a high-risk phenotype for AUD through altered alcohol behavioral sensitivity. Our study highlights the importance of future studies that seek to identify the neurobiological mechanisms underlying these interactions, which may pave the way for targeted interventions in populations affected by childhood adversity and excessive alcohol consumption.

## INTRODUCTION

Adverse childhood experiences (ACEs), such as abuse or neglect, are powerful risk factors for developing alcohol use disorder (AUD) in adulthood^1-3^. In the United States, more than 60% of adults report experiencing at least one type of ACE, and approximately 17% report experiencing four or more types of ACEs, with women and people of low socioeconomic status most likely to report a higher number of ACEs^4,5^. In adolescence, female high school students are more likely to drink alcohol and be diagnosed with AUD than their male peers^6-8^. In adulthood, sex differences in alcohol use and abuse reverse, with men consistently more likely to drink alcohol, engage in binge and high-intensity drinking, and be diagnosed with AUD^8-12^. Collectively, these data highlight the complex interactions between childhood adversity, gender/biological sex, and the development of AUD in adulthood, underscoring the importance of causally investigating how ACEs modulate AUD susceptibility.

Notably, alcohol produces dose-dependent biphasic effects of alcohol in both humans and mice – stimulatory at low doses, sedative at high doses^13^ – and altered behavioral sensitivity to alcohol is strongly linked to AUD susceptibility in humans^14-16^. Abundant clinical evidence points to complex interactions between stress and alcohol drinking^17^, but little is known about how biological sex and ACEs interact to modulate behavioral sensitivity to alcohol, and in turn, AUD susceptibility. Given the limitations of disentangling these influences in humans, mouse models of early life stress (ELS) have been developed using a variety of methods, including low bedding/nesting (LBN) and/or maternal separation (MS) paradigms. These paradigms are generally imposed during the first two postnatal weeks and are designed to experimentally produce adverse maternal care, including rough handling, increased time away from the nest, and reduced nursing and grooming^18-20^.

In C57BL/6J mice, foundational work investigating sex differences in behavioral sensitivity to alcohol generally points to heightened sensitivity in males relative to females. Early studies reported that males displayed greater locomotor activity in response to low doses of alcohol^21^ and longer loss of righting reflex duration following high doses of alcohol^22^, as well as greater impairment in some measures of motor coordination following medium doses of alcohol^23^. A recent study reported increased sensitivity to both the stimulatory and sedative effects of alcohol following unpredictable, chronic, mild stress in adult males, but not females, suggesting sex-specific modulation of stress effects on behavioral sensitivity to alcohol^24^. Given the converging lines evidence in mice and humans that 1) childhood adversity/early life stress confers increased risk of AUD susceptibility, 2) alcohol behavioral sensitivity influences AUD susceptibility, and 3) stress modulates alcohol behavioral sensitivity, we hypothesized that ELS modulates behavioral sensitivity to alcohol, and in turn, promotes escalation of alcohol drinking in a sex-specific manner.

There are currently no published studies in C57BL/6J background mice examining the sex-specific effects of ELS on alcohol behavioral sensitivity and long-term intermittent-access voluntary alcohol drinking. Existing related studies have reported variable effects of ELS on alcohol drinking and other AUD-associated behaviors, including anxiety, depression, and social behaviors, likely due to differences in ELS paradigms, alcohol drinking models, developmental age at testing, and limited examinations of sex differences^25-27^. Therefore, we sought to resolve outstanding gaps in the literature specifically regarding the examination of: 1) a combined LBN+MS ELS paradigm, 2) sex differences, 3) behavioral effects in adulthood rather than adolescence, 4) behavioral sensitivity to both stimulatory and sedative doses of alcohol, 5) voluntary alcohol drinking in the absence of forced dependence, and 6) interactive effects of ELS and alcohol drinking on sociability, anxiety, and other motivated behaviors.

Here, we implemented a robust LBN+MS model to systematically examine how ELS modulates 1) behavioral sensitivity to both the stimulatory and sedative effects of alcohol (locomotor stimulation, loss of righting reflex), 2) escalation of voluntary alcohol drinking (long-term two-bottle choice intermittent access), and 3) social and affective behaviors following alcohol drinking (sociability, social novelty preference, anxiety-like behaviors, sucrose preference) in adult female and male C57BL/6J background mice. Our results indicate that ELS generates a high-risk phenotype wherein ELS mice demonstrate greater locomotor stimulation and diminished sedation in response to alcohol, enhanced alcohol consumption and preference over time, and social deficits produced by combined ELS and alcohol drinking experience. We also identified important sex differences across distinct impacts of ELS and tracked estrous cycles in female mice to evaluate potential influences of hormonal fluctuations. Overall, our findings suggest the utility of the LBN+MS ELS model for future investigations to uncover the neurocircuit basis of ELS/ACE impacts on AUD-like susceptibility, ultimately advancing precision therapeutic strategies for treating psychiatric conditions of stress and addiction.

## METHODS

### Mice

Homozygous CRF-ires-Cre males (B6(Cg)-Crh^tm1(cre)Zjh^/J), JAX strain # 012704) and B6 females (C57BL/6J, JAX strain #000664) were purchased and bred in house. To generate experimental litters, two experienced B6 female breeders (3-6 months old, previously gave birth to and successfully weaned at least one litter containing 5 or more pups) were trio bred with one CRF-Cre^+/+^ male mouse (2-12 months old). Approximately 16 days after pairing, dams were assessed for pregnancy and separated into their own new, clean cages. Pregnant dams were checked twice daily for the birth of pups (P0). Litters were composed of heterozygous CRF-Cre^+/-^ females and males ranging from 5-12 pups per litter. Mice were weaned, ear punched, and weighed at P25 and housed with same sex-littermates in groups of 2-5 mice. Body weights were measured regularly until adulthood at P28, P42, P56, and P70. Breeders and weaned mice were housed in ventilated Innovive cages containing adequate bedding and nesting material (Envrio-dri and one cotton nestlet) but no enrichment objects (paper tubes, etc.) Mice were maintained on a 12 hr light/dark cycle (lights on 07:00) in a temperature- and humidity-controlled facility with ad libitum access to standard chow (Teklad Irradiated Global 18% Protein Rodent Diet, product # 2918) and water. Two weeks prior to behavioral testing, adult mice (>P56) were transferred to standard static mouse cages and housed in a reverse 12 hr light/dark cycle room (lights on 19:00) in the same facility. Cages were randomly distributed on the rack shelves to minimize confounding effects of rack placement^28^. Mice remained group-housed with ad libitum access to standard chow and water, unless otherwise specified. All behavioral tests were conducted in adult (>P70) female and male mice. All animal procedures were approved and conducted in accordance with the Institutional Animal Care and Use Committee of Stanford University.

### Early Life Stress

Litters from paired dams were randomly assigned to undergo either control (CTL) or early life stress (ELS) rearing conditions. CTL litters experienced sufficient bedding and nesting conditions and uninterrupted maternal care from birth to weaning (P25). ELS litters experienced both limited bedding and nesting (LBN) conditions and repeated maternal separation (MS) from P2-P12. For LBN, approximately 75% of bedding and nesting materials (Enviro-dri and cotton nestlet) were removed from the cage on P2 and remained limited 24 hr/day until P12. For MS, the dam was removed from the home cage for 4 hrs/day (from approximately 9 AM - 1 PM) and housed in a separate cage with ad libitum food and water while pups remained together in the home cage. Following the final round of MS on P12, ELS litters and dams were housed in a new, clean cage with adequate nesting and bedding material until weaning (P25). CTL litters were also given a cage change on P12. Dams assigned to CTL conditions were used as breeders for additional litters if <6 months old at weaning. Dams assigned to ELS conditions were euthanized following weaning, regardless of their age.

### Maternal Nest Entries

On P2, P3, P4, P8, P12, ELS dams were videorecorded in their home cage for 30 minutes following reunion from maternal separation. CTL dams were briefly lifted and replaced into their home cage and recorded for 30 minutes at a similar time of day. Nest entries, classified as at least half of the dam’s body entering the area where pups were laying, were hand scored using BORIS software by a single experimenter.

### Locomotor Activity (Stimulation Sensitivity)

Mice were moved to a red-light behavior room approximately two hours into their dark cycle and left to acclimate for one hour. On Day 1, mice received an intraperitoneal (i.p.) injection of 0.25 ml sterile saline (Sal) and were immediately placed into an empty standard static mouse cage. Their activity was videorecorded for 20 minutes, after which they were returned to their home cage and replaced in their colony room. On Day 2, mice received an i.p. injection of a stimulatory dose (1.75 g/kg) of 20% ethanol (EtOH) in sterile saline and were immediately placed into an empty static cage to record activity for 20 minutes. Mice received alternating days of saline and ethanol injections for a total of 8 days (4 Sal and 4 EtOH). Distance traveled was quantified using ezTrack software and converted to average velocity (cm/s).

### Loss of Righting Reflex (Sedation Sensitivity)

Mice were moved to a red-light behavior room approximately two hours into their dark cycle and left to acclimate for one hour. Each mouse was placed in an empty static cage for one minute before receiving an i.p. injection of a sedative dose (4.0 g/kg) of 20% ethanol (EtOH) in sterile saline, then immediately placed back in the empty cage. After 45 seconds, mice were turned on their side every 10 seconds to check for loss of righting reflex (LORR), or the inability to right themselves onto all four paws. Mice were continuously monitored for recovery of the righting reflex. Specifically, once mice righted themselves onto all four paws, they were turned back on their side three more times, and if able to right themselves three times within 30 seconds, their righting reflex was considered regained. The times of injection, loss of righting reflex, and recovery of righting reflex were recorded and used to calculate latency to LORR (injection to loss) and duration of LORR (loss to recovery).

### Long-Term Two-Bottle Choice Intermittent Access Alcohol Drinking

Mice were singly housed in standard static mouse cages with two 25 ml graduated cylinders fitted with sippers as water bottles and transferred to a reverse light cycle room (described above). Following two weeks of acclimation, mice underwent a week of baseline measurements (Week 0) during which their daily water intake, food intake, and body weight was recorded. For the next eight weeks (Weeks 1-8), mice underwent two-bottle choice intermittent access (2BC-IA) alcohol drinking. On Sunday, Tuesday, and Thursday of each week, one water bottle was removed and replaced with a bottle containing 20% v/v ethanol in water approximately three hours into the dark cycle (ON). Alcohol bottle placement was counterbalanced based on side preference determined during Week 0, so that each group had a baseline “alcohol-side” preference between 45-55%. Ethanol bottles were removed after 24 hours and replaced with a water bottle (OFF), with access to only water for the following 24 hours. Fluid levels in each bottle and food weights were recorded daily at ON and OFF times, and body weights were recorded weekly on Sundays. Mice always maintained ad libitum access to chow.

### Short-Term Two-Bottle Choice Limited Access Alcohol Drinking in the Dark

As for chronic alcohol drinking, mice were singly housed in standard static mouse cages with two 25 ml graduated cylinders fitted with sippers as water bottles and transferred to a reverse light cycle room (described above). Following two weeks of acclimation, mice underwent a week of baseline measurements (Week 0) during which their daily body weight and water and food intake were recorded. For the next four weeks (Week 1-4), mice underwent four days per week of two-bottle choice limited access (2BC-LA) alcohol drinking in the dark (DID). From Monday-Thursday, approximately three hours into the dark cycle, one water bottle was removed and replaced with a bottle containing 20% v/v and a single piece of chow was placed on the floor of the cage (ON). After four hours of alcohol access, the ethanol bottle was removed and replaced with a water bottle, and all chow was returned to the food hopper (OFF). Fluid levels in each bottle and food weights were recorded daily at ON and OFF times, and body weights were recorded weekly on Mondays.

### Short-Term Two-Bottle Choice Limited Access Sucrose Drinking in the Dark

Following a week of alcohol abstinence (Week 5), mice underwent two weeks (Weeks 6-7) of 2BC-LA sucrose DID. All procedures were the same as described for alcohol DID, except one water bottle was replaced with a bottle containing 0.5% w/v sucrose in water. Sucrose bottles were placed on the same side as ethanol bottles during alcohol DID.

### Anxiety-Like Behavior Tests

All mice tested for anxiety-like behavior were singly housed in standard static mouse cages in a reverse light cycle room (described above). Tests were conducted in red light approximately 3-6 hours into the dark cycle on Monday (EPM), Wednesday (OFT), and Friday (NSFT) of the same week. For naïve anxiety tests, EPM, OFT, and NSFT were conducted in the home colony room following two weeks of reverse light cycle single housing acclimation. For post-EtOH drinking anxiety tests, EPM, OFT, and NSFT were conducted in the mice’s home colony room following four weeks of alcohol DID, 4, 6 or, 8 days after their final drinking session, respectively. Only female experimenters were present in the room during all anxiety-like behavior testing. EzTrack was used to quantify total distance traveled and time spent in open and closed arms (EPM), center (OFT), and food zone (NSFT).

*Elevated Plus Maze (EPM):* The EPM apparatus (San Diego Instruments, product # 7001-0336) consisted of two sets of perpendicular arms, two open and two enclosed (each 30 x 5 cm) with a center connecting square (5 x 5 cm), elevated 39 cm above the floor. Mice were placed in the center square facing one of the open arms and videorecorded for 5 minutes.

*Open Field Test (OFT):* The open field apparatus (San Diego Instruments, product # 7001-0354) consisted of four square arenas (each 50 x 50 cm), with the “center zone” defined as the inner 25 x 25 cm square. Mice were placed in the center of the arena and video recorded for 5 minutes.

*Novelty-Suppressed Feeding Test (NSFT):* Mice were food deprived for 24 hours prior to the test. A chow pellet was placed in the center of the open field arena (described above) and held in place using a small amount of mounting putty (Scotch, product # 860). The “food zone” was defined as the innermost 12.5 x 12.5 cm square of the arena immediately surrounding the chow. Mice were placed in the outermost corner of the arena and videorecorded for 10 minutes. After the test, mice were returned to their home cage and the piece of chow was placed on the floor of the cage. Pellets were weighed before and after the 10 minute test, as well as at two and four hours after mice were returned to their home cage.

### Pre-Social Behavior Water or Ethanol Drinking

Mice were singly housed in standard static mouse cages with two 25 ml graduated cylinders fitted with sippers as water bottles and transferred to a reverse light cycle room (described above). Following two weeks of acclimation, mice underwent a week of baseline measurements (Week 0) during which their daily water intake, food intake, and body weight was recorded. For next 3 weeks (Week 1-3), mice were randomly split into H2O or EtOH groups. Mice in the H2O group received 24/7 access to two water bottles, while mice in the EtOH group underwent 2BC-IA alcohol drinking (described above).

### Social Behavior Tests

Four days after the final alcohol access day, mice were tested for sociability and social novelty using the three-chamber social behavior test, as described^29,30^. On the day prior to testing, test mice were habituated to the apparatus (constructed in-house, similar to San Diego Instruments, product # 7500-0362) for 10 minutes, and stimuli mice (male and female 4-7 week old C57BL/6J mice) were habituated to being placed under wire mesh cups for 10 minutes.

On the day of testing, mice were moved to a red-light behavior room approximately two hours into their dark cycle and left to acclimate for one hour. Test mice were placed in the center holding chamber of the apparatus and wire mesh cups were placed upside down in the other two chambers. To begin each test phase, the transparent chamber dividers were removed, and the test mouse was videorecorded for 10 minutes, after which it was re-confined to the center holding chamber. In phase 1 (Habituation), both cups remained empty. In phase 2 (Sociability), one cup contained a novel mouse (same-sex juvenile) and the other contained a novel object (black plastic mouse). In phase 3 (Social Novelty), one cup contained a familiar mouse (the novel same-sex juvenile from phase 2) and the other contained novel mouse (a new same-sex juvenile). Videos were hand scored by a single experimenter for interaction time with each stimulus, defined as sniffing, climbing, and/or repeated circling of the cup.

### Estrous Staging

Female mice were estrous staged via vaginal cytology as described^31,32^. Briefly, cells were collected from the vaginal canal via lavage with sterile double distilled water. Collected fluid was placed on a glass slide and proportions of vaginal cell types were examined via light microscopy. Samples containing > 50% leukocytes were staged as low estradiol/E2 (metestrus/diestrus), and samples containing > 50% nucleated or cornified squamous epithelial cells were staged as high estradiol/E2 (proestrus/estrus). Samples not containing enough cells to definitively stage are marked as unstaged. Estrous cycle length represents the number of days from the first day of high E2 to the final day of low E2 before returning to high E2. Range of estrous cycle length represents the difference between and individual mouse’s longest and shortest estrous cycle lengths. Vaginal lavage was performed following behavior testing (alcohol stimulation and sensitivity tests) or before/after alcohol bottle access on ON/OFF ethanol drinking days, respectively. Male mice received control handling for 20 seconds during the same time period.

### Data Analysis

Experimenters were blind to ELS condition (but not sex, due to estrous staging or need to pair with same-sex social stimulus) during behavior testing, and were blind to all relevant factors (ELS, Sex, EtOH) during video/data analysis. Raw data from females and males were analyzed separately for all experiments. Statistical tests were performed in GraphPad Prism 10. Significance threshold was set at p < 0.05. On figures, significant main effects are noted by * (p < 0.05), ** (p < 0.01), *** (p < 0.001), and **** (p < 0.0001) and data are expressed as mean +/- standard error of the mean (SEM). Figures were created with GraphPad Prism 10 and Biorender.

### Maternal Nest Entries

The number of nest entries were analyzed by 2-way RM ANOVA with factors of ELS (CTL or ELS) and Postnatal Day. Sidak’s multiple comparisons tests were conducted post-hoc.

### Body Weights

Body weight at weaning was analyzed by 2-way ANOVA with factors of ELS (CTL or ELS) and Sex (Female or Male). Biweekly bodyweights were analyzed by 3-way RM ANOVA with factors of ELS (CTL or ELS), Sex (Female or Male), and Age.

### EtOH Stimulation

Velocity was analyzed by 3-way RM ANOVA with factors of ELS (CTL or ELS), EtOH (saline or EtOH injection), and Pair (Sal-EtOH pairings). ΔVelocity was analyzed for each Sal-EtOH pair by 2-way RM ANOVA with factors of ELS (CTL or ELS) and Pair (Sal-EtOH pairings). Average ΔVelocity was analyzed by unpaired t test.

### EtOH Sedation

Latency and duration were analyzed by mixed-effects analysis with factors of ELS (CTL or ELS) and Day. 2-way RM ANOVA could not be used due to missing values, as all mice did not lose righting reflex on all three days of testing. Average latency and duration were analyzed by unpaired t-test.

### EtOH/Sucrose/H2O Drinking

Raw values of daily water (ml), ethanol (ml), and food (g) consumption were used to identify outliers +/- 2.5 standard deviations from the mean of each group across all 8 weeks of data collection. Outliers were excluded from all further analysis. Daily ethanol consumption (ml) was converted to grams based on density of EtOH concentration and divided by the animal’s body weight to give daily ethanol intake expressed as g/kg. Daily water, total liquid (ethanol + water), and food consumption were divided by the animal’s body weight to give daily water and total liquid intake expressed as ml/kg and daily food intake expressed as g/kg. Daily ethanol preference was calculated by dividing total ethanol consumption (ml) by total liquid consumption (ml) and converting to a percentage. Values were averaged across three (24hr 2BC-IA) or four (4hr DID) drinking sessions per week for each animal. If two or more daily values in a week were missing or excluded for an animal, a weekly average was not calculated for that week. Weekly averages were analyzed by 2-way RM ANOVA or mixed effects analysis with factors of ELS (CTL or ELS) and Week. Sidak’s multiple comparisons tests were conducted post-hoc. Differences between first and last week averages were analyzed by unpaired t-test. For pre-social behavior drinking, weekly averages for total liquid and food intake were analyzed by 3-way RM ANOVA with factors of ELS (CTL or ELS), EtOH (H2O or EtOH), and Week and differences between first and last week averages were analyzed by 2-way ANOVA with factors of ELS (CTL or ELS) and EtOH (H2O or EtOH).

### Anxiety-Like Behavior

Distance traveled (cm) was divided by total test time (s) to give average velocity expressed as cm/s. Percent open arm (EPM), center (OFT), and food zone (NSFT) were calculated by dividing the time spent in the region of interest (s) by total test time (s) and converting to a percentage. Values were analyzed by 2-way ANOVA with factors of ELS (CTL or ELS) and EtOH (Naïve or EtOH).

### Social Behavior

Raw values of interaction times were used to identify and exclude outliers +/- 2.5 standard deviations from the mean of all mice interacting with each stimulus (novel mouse #1, novel object, familiar mouse, novel mouse #2). Interaction time was analyzed by 3-way ANOVA with factors of ELS (CTL or ELS), EtOH (H2O or EtOH), and Stimulus (Mouse or Object; Familiar or Novel). Sidak’s multiple comparisons tests were conducted post-hoc.

### Estrous

Average cycle length and range of cycle length were analyzed by unpaired t-test. For EtOH stimulation, saline and EtOH velocities were averaged across low and high E2 stages for each mouse. For EtOH sedation, latency and duration for low and high E2 represent either a single data point (one day) or an average of data points (two days) for each mouse. Mice were excluded from estrous analysis if they were not staged in low and high E2 for at least one day each across three days of tests. For EtOH drinking, intake values were averaged across low and high E2 stages during Weeks 1-4 (12 total drinking sessions) for each mouse. Mice were excluded from estrous analysis if they were not staged in low and high E2 for at least three days each across four weeks of drinking. Values were analyzed by 2-way RM ANOVA with factors of Estrous (LowE2 or HighE2) and ELS (CTL or ELS).

## RESULTS

### ELS rearing conditions induce fragmented maternal care and decrease offspring body weights

To model ELS, matched litters were randomly assigned to undergo normal (CTL) or combined limited bedding/nesting and repeated maternal separation (ELS) rearing conditions (Fig. 1A). There was no difference in the number of pups per litter in CTL and ELS conditions (Fig. 1B). To validate the LBN+MS ELS model, we assessed phenotypes previously reported in LBN and/or MS models, including increased maternal nest entries and exits and decreased body weight of pups at weaning^20,33-35^. ELS dams displayed significantly more nest entries/exits following reunion from maternal separation relative to CTL dams, particularly on the first day (P2) of separation (Fig. 1C), indicating increased fragmentation of maternal care. At weaning, both female and male ELS mice weighed significantly less than same-sex CTL mice (Fig. 1D). Mice were weighed bi-weekly at P28, P42, P56, and P70 to determine whether this difference persisted into adulthood, and we found that both female and male ELS mice continued to weigh significantly less than same-sex CTL mice throughout this period (Fig. 1E). Taken together, these results validated the ability of our combined LBN+MS ELS model to generate expected adverse maternal care and physiological impacts in offspring.

**Figure 1:**
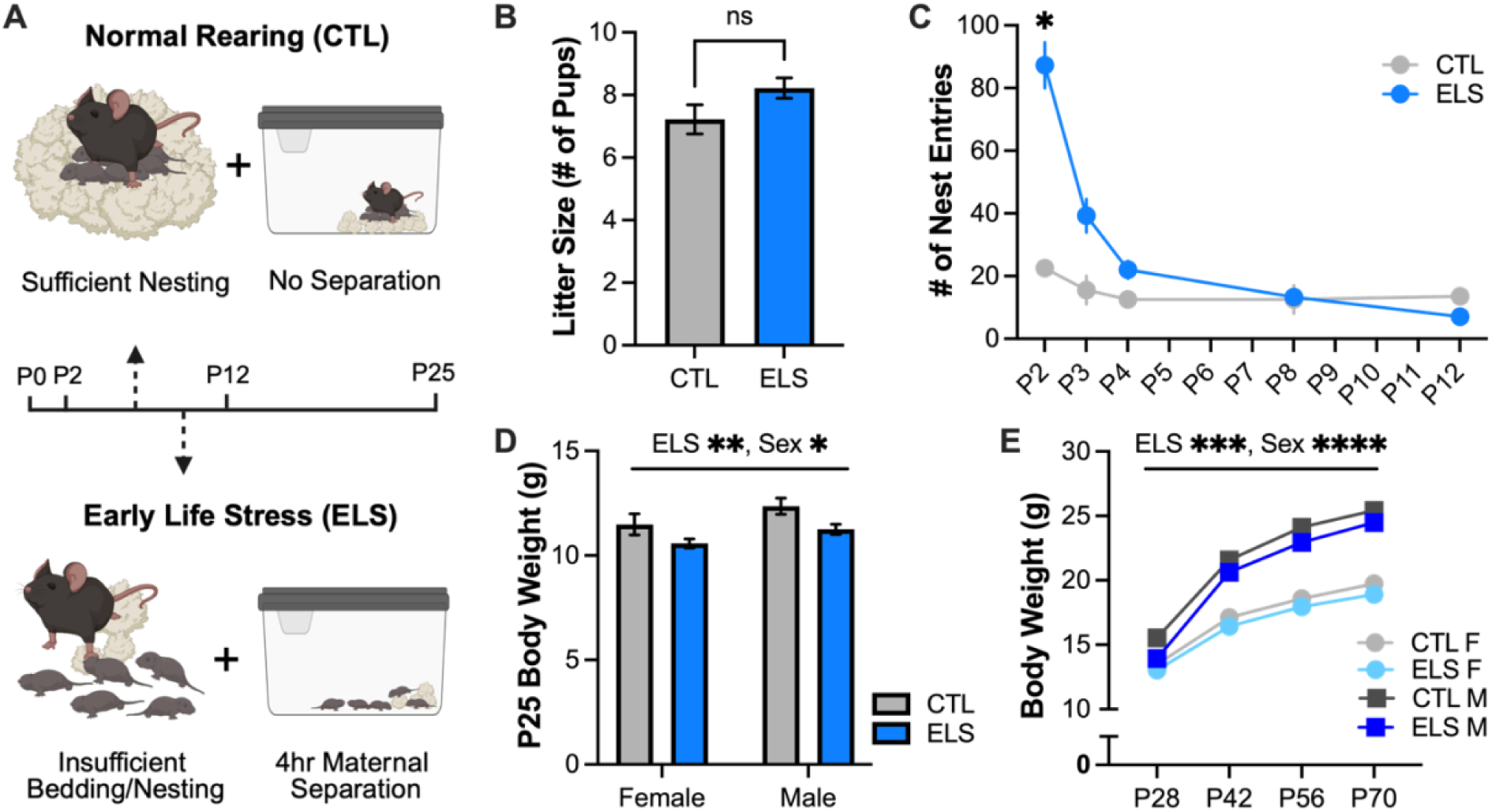
Validation of LBN+MS ELS model. **A** Schematic of CTL and ELS rearing conditions. **B** Number of pups in litters assigned to CTL and ELS. No difference between groups (p = 0.0966, t^16^ = 1.765) **C** Number of maternal nest entries in a 30-minute period following maternal reunion (ELS) or at a similar time of day (CTL). Main effects of *ELS (p = 0.0125, F^1, 4^ = 18.61), ****Postnatal Day (p < 0.0001, F^1.994, 7.974^ = 36.77), and an ****ELS x Postnatal Day interaction (p < 0.0001, F^4, 16^ = 22.17). Significant difference between CTL and ELS dams on *P2 (p = 0.0145) **D** Body weight of female and male CTL and ELS mice at weaning on P25. Main effects of **ELS (p = 0.0033, F^1, 120^ = 8.984) and *Sex (p = 0.0229, F^1, 120^ = 5.310). **E** Body weights at 4, 6, 8 and 10 weeks old. Main effects of ***ELS (p = 0.0006, F^1, 95^ = 12.75), ****Sex (p < 0.0001, F^1, 95^ = 267.0), and ****Week (p < 0.0001, F^2.326, 220.9^ = 2064). B: n = 9 litters/condition C: n = 2 CTL dams, 4 ELS dams D: n = 25 CTL F, 28 ELS F, 30 CTL M, 39 ELS M E: n = 22 CTL F, 24 ELS F, 24 CTL M, 29 ELS M

### ELS increases behavioral sensitivity to the stimulant effects of alcohol in females

Given the established link between ACEs/ELS and predisposition for AUD, we hypothesized that ELS may increase and decrease sensitivity to the stimulatory (appetitive) and sedative (aversive) effects of alcohol, respectively. To investigate the effects of ELS on alcohol stimulation sensitivity, mice received intraperitoneal (i.p.) injections of saline or low dose EtOH (1.75 g/kg) during their active phase on alternating days for four saline/EtOH pairings across eight consecutive days and assessed for locomotor activity (Fig. 2A). In females, we found that EtOH modulated locomotor activity in ELS but not CTL mice (Fig. 2B). To examine individual differences in locomotor activity across days, we calculated the difference in velocity for each pairing (EtOH minus saline; EtOH-Sal). ELS females displayed increased activity in response to EtOH relative to saline, while CTL females displayed no difference in EtOH-Sal velocity (Fig. 2C). ELS significantly increased the difference in EtOH-Sal locomotor activity when analyzed across repeated pairings and averaged across all pairings (Fig. 2C inset). In males, neither ELS nor EtOH modulated locomotor activity (Fig. 2D), and ELS did not significantly affect the difference in EtOH-Sal velocity across pairings (Fig. 2E).

**Figure 2:**
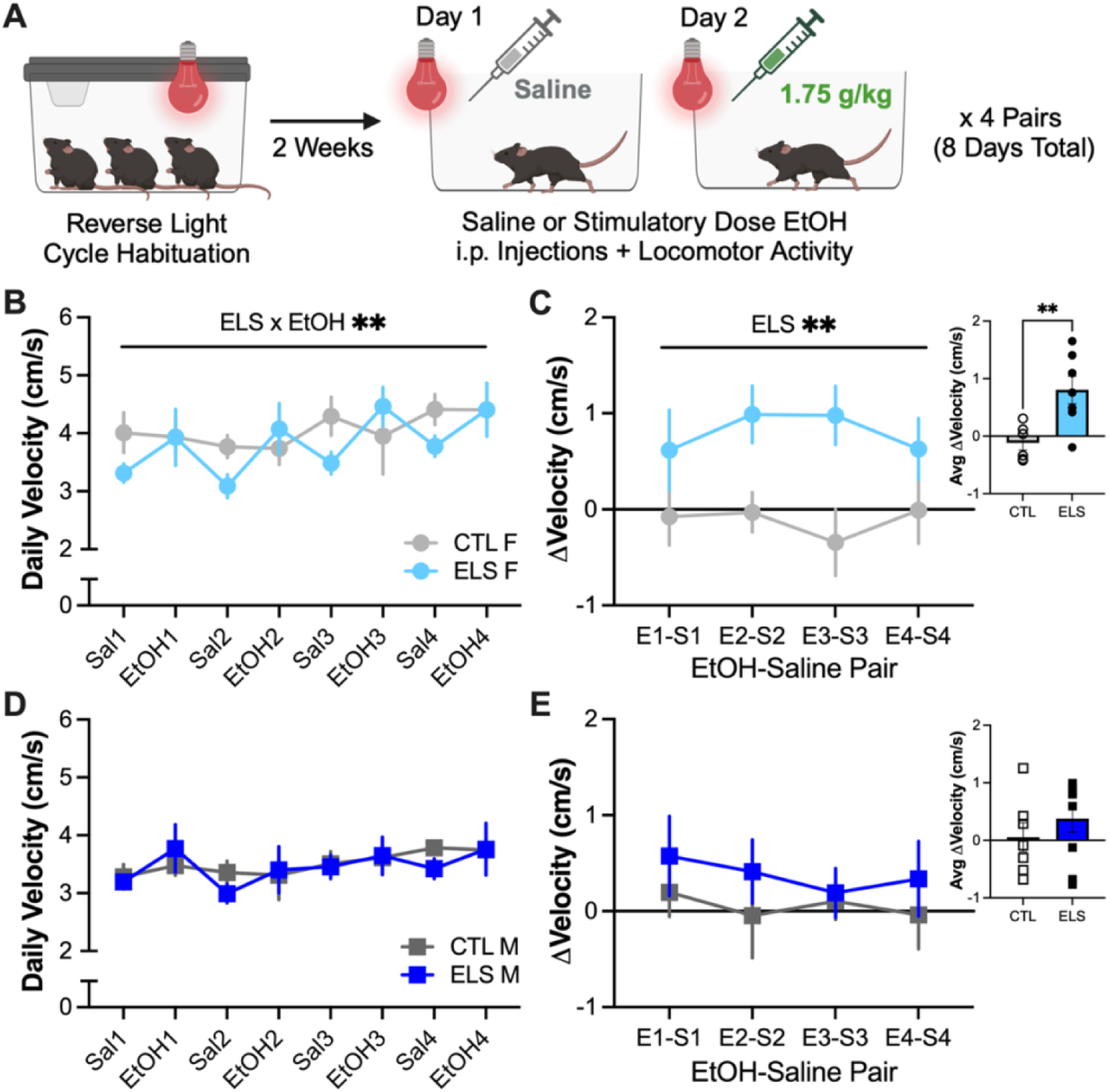
ELS increases behavioral sensitivity to the stimulant effects of alcohol in females. **A** Schematic of stimulatory alcohol sensitivity experimental design. **B** Velocity following Sal or EtOH injection in CTL and ELS females. Main effects of *EtOH (p = 0.0343, F^1, 11^ = 5.831), *Pair (p = 0.0164, F^3, 33^ = 3.948), and an **ELS x EtOH interaction (p = 0.0081, F^1, 11^ = 10.39). **C** V^EtOH^ – V^Sal^ for each Sal-EtOH pair in CTL and ELS females. Main effect of **ELS (p = 0.0081, F^1, 11^ = 10.39). Inset: Average V^EtOH^ – V^Sal^ across 4 Sal-EtOH pairs. **Significant difference between groups (p = 0.0081, t^11^ = 3.224). **D** Velocity following i.p. saline Sal or EtOH injection in CTL and ELS males. No main effect of ELS, EtOH, or Pair. **E** V^EtOH^ – V^Sal^ for each Sal-EtOH pair in CTL and ELS males. No main effect of ELS or Pair. Inset: Average V^EtOH^ – V^Sal^ across 4 Sal-EtOH pairs. No significant difference between groups. n = 6 CTL F, 7 ELS F, 7 CTL M, 9 ELS M

### ELS decreases behavioral sensitivity to the sedative effects of alcohol, particularly in males

To investigate the effects of ELS on alcohol sedation sensitivity, mice received i.p. injections of high dose EtOH (4.0 g/kg) during their active phase on three consecutive days and assessed for loss of righting reflex (LORR) (Fig. 3A). In females, ELS significantly increased LORR latency when analyzed across repeated days (Fig. 3B) but did not affect latency when averaged across days (Fig. 3B inset). Similarly, there was a trend towards decreased LORR duration in ELS females when analyzed across repeated days (Fig. 3C), but no significant effect of ELS on duration when averaged across days (Fig. 3C inset). In males, ELS did not affect LORR latency (Fig. 3D), but did significantly decrease LORR duration (Fig. 3E). Taken together, the enhanced sensitivity to the stimulant effects of alcohol (Fig. 2), and reduced sensitivity to the sedative effects of alcohol (Fig. 3), in ELS mice parallels the pattern of altered alcohol sensitivity in humans with heightened predisposition to AUD.

**Figure 3:**
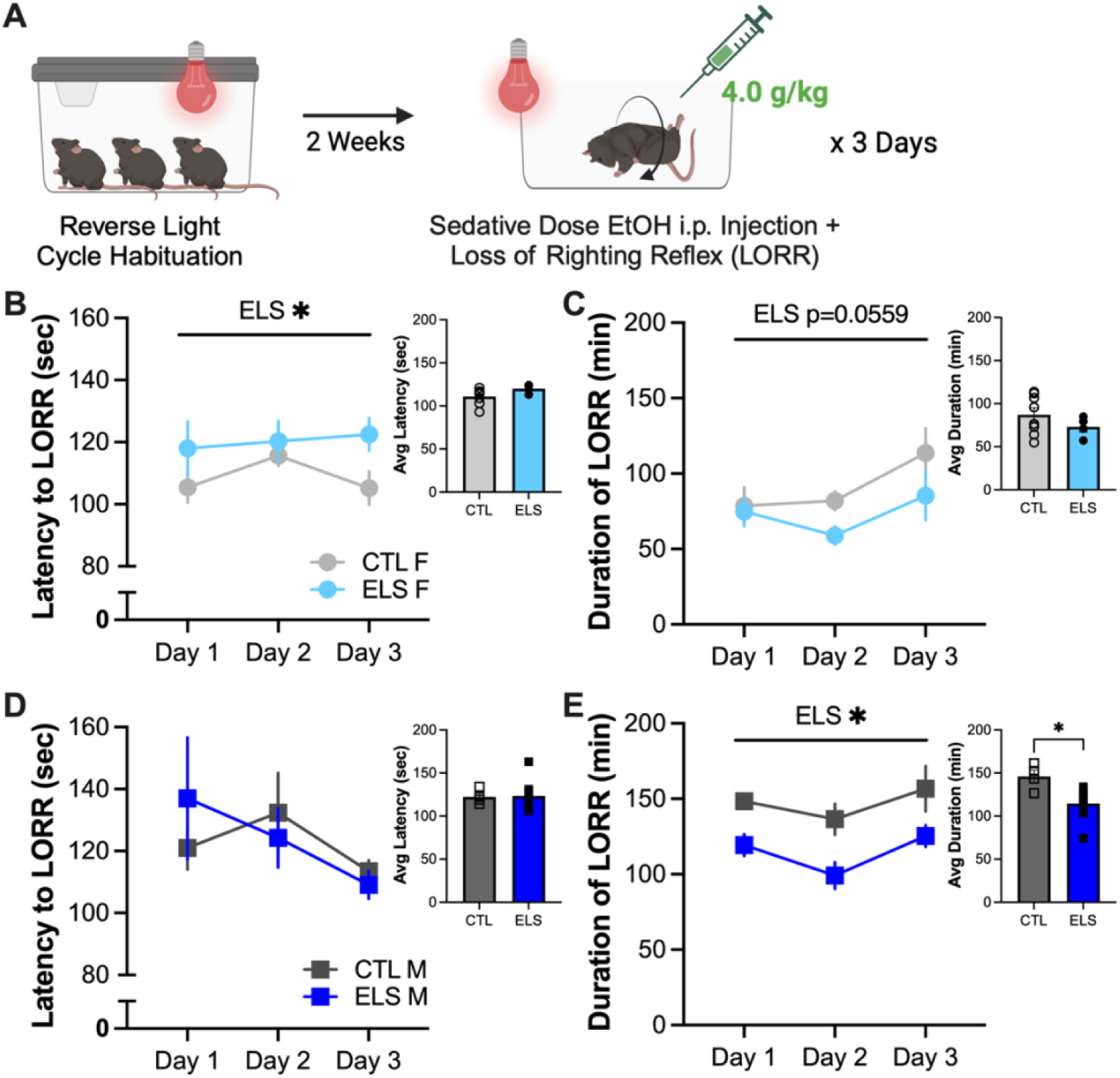
ELS decreases behavioral sensitivity to the sedative effects of alcohol, particularly in males. **A** Schematic of sedative alcohol sensitivity experimental design. **B** Latency to lose righting reflex in CTL and ELS females. Main effect of *ELS (p = 0.0360, F^1, 10^ = 5.860). Inset: Average latency to lose righting reflex across three days. No difference between groups. **C** Duration of loss of righting reflex in CTL and ELS females. Trend towards a main effect of Day (p = 0.0559, F^1.435, 11.48^ = 4.094). Inset: Average duration of loss of righting reflex across three days. No difference between groups. **D** Latency to lose righting reflex in CTL and ELS males. No main effect of ELS or Day. Inset: Average latency to lose righting reflex across three days. No difference between groups. **E** Duration of loss of righting reflex in CTL and ELS males. Main effects of *ELS (p = 0.0264, F^1, 10^ = 6.773) and *Day (p = 0.0106, F^1.951, 15.61^ = 6.226). Inset: Average duration of loss of righting reflex across three days. *Significant difference between groups (p = 0.0156, t = 2.910, df = 10). n = 8 CTL F, 4 ELS F, 3 CTL M, 8 ELS M

### ELS increases voluntary alcohol intake and preference in a long-term drinking model, particularly in males

Given that ELS enhanced sensitivity to stimulant effects of alcohol and reduced sensitivity to sedative effects of alcohol, we next hypothesized that ELS might also drive increased escalation of voluntary alcohol drinking. To investigate the effects of ELS on alcohol drinking, we used a long-term two-bottle choice intermittent access (2BC-IA) alcohol drinking model with three 24-hour drinking sessions per week over eight weeks (Fig. 4A).

**Figure 4:**
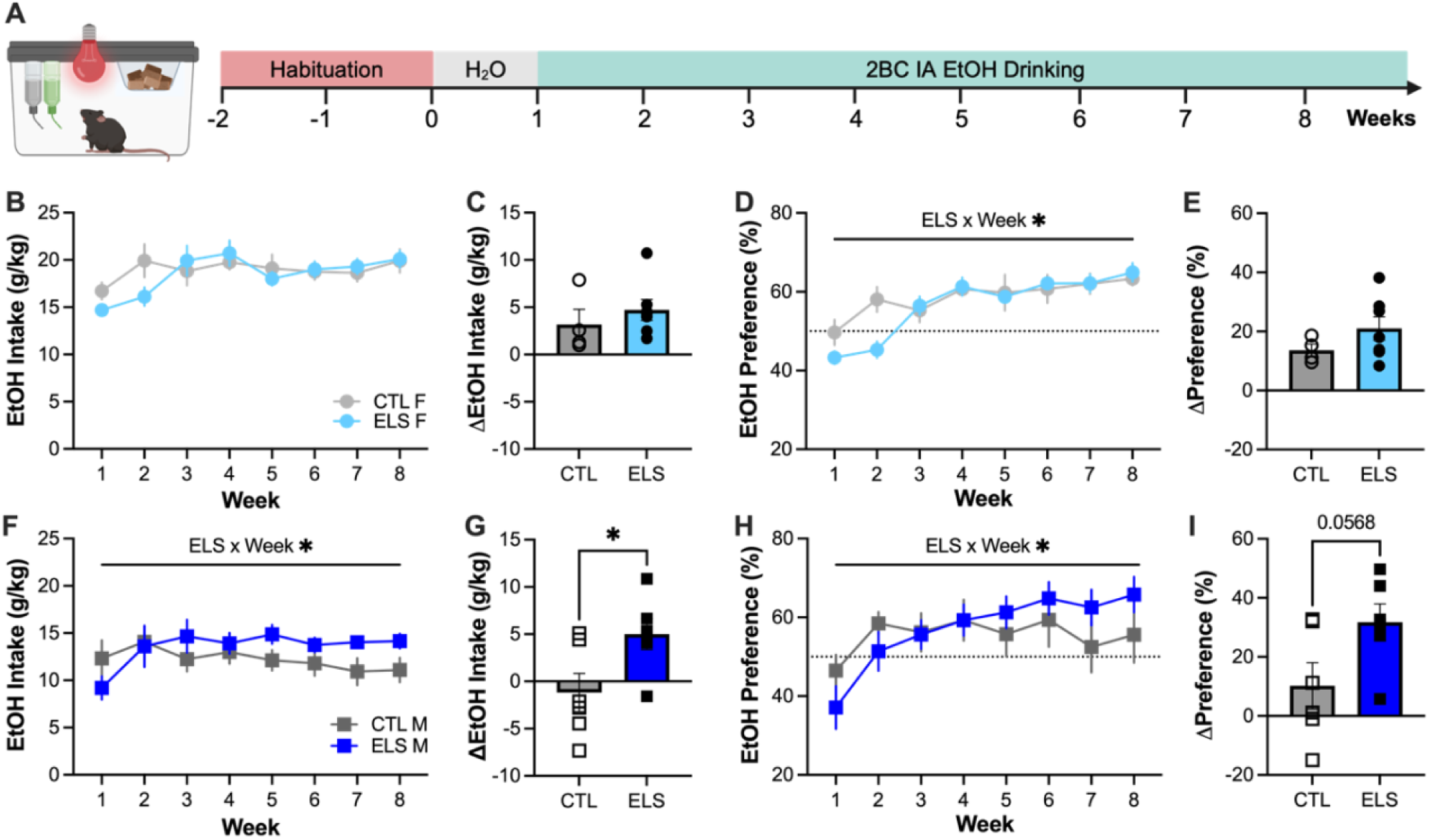
ELS increases long-term alcohol drinking and preference, particularly in males. **A** Schematic of long-term 2BC-IA alcohol drinking. **B** Ethanol intake in CTL and ELS females. Main effect of **Week (p = 0.0057, F^4.047, 38.16^ = 4.280). **C** Difference in ethanol intake between Week 8 and Week 1 in CTL and ELS females. No difference between groups. **D** Ethanol preference in CTL and ELS females. Main effects of ****Week (p < 0.0001, F^4.212, 39.72^ = 13.68) and *ELS x Week interaction (p = 0.0289, F^7, 66^ = 2.417). **E** Difference in ethanol preference between Week 8 and Week 1 in CTL and ELS females. No difference between groups. **F** Ethanol intake in CTL and ELS males. Main effect of *ELS x Week interaction (p = 0.0497, F^7, 90^ = 2.116). **G** Difference in ethanol intake between Week 8 and Week 1 in CTL and ELS males. *Significant difference between groups (p = 0.0388, t = 2.377, df = 10). **H** Ethanol preference in CTL and ELS males. Main effects of ***Week (p = 0.0003, F^2.677, 34.42^ = 8.763) and *ELS x Week interaction (p = 0.0238, F^7, 90^ = 2.454). **I** Difference in ethanol preference between Week 8 and Week 1 in CTL and ELS males. Trend towards a difference between groups (p = 0.0568, t = 2.152, df = 10). n = 4 CTL F, 8 ELS F, 8 CTL M, 8 ELS M

In females, ELS did not significantly affect weekly EtOH intake (Fig. 4B) or escalation of EtOH intake from Week 1 to Week 8 (Fig. 4C). Relative to CTL females, ELS females displayed a significantly different pattern of EtOH preference across weeks (Fig. 4D), however, there was no significant effect of ELS on escalation of EtOH preference from Week 1 to Week 8 (Fig. 4E). In males, ELS mice displayed a significantly different pattern of EtOH drinking across weeks (Fig. 4F) and significantly greater escalation of EtOH intake relative to CTL mice (Fig. 4G). Similarly, ELS males displayed a significantly different pattern of EtOH preference across weeks relative to CTL males (Fig. 4H) and trended towards increased escalation of EtOH preference (Fig. 4I).

In both females and males, we did not identify any significant effects of ELS on escalation of water intake, total liquid intake, or food intake during chronic 2BC-IA alcohol drinking (Supplemental Fig. 1A-F), indicating that the effects of ELS on EtOH intake and preference are not likely explained by altered fluid balance or caloric needs.

We also investigated the effects of ELS on alcohol drinking using a two-bottle choice limited access (2BC-LA) alcohol drinking in the dark (DID) model with four 4-hour drinking sessions per week over four weeks. We did not identify any significant effects of ELS in females or males on weekly EtOH intake, EtOH preference, water intake, total liquid intake, or food intake, nor on escalation of these measures between Week 1 and Week 4 (Supplemental Fig. 2A-J), suggesting that the limited access DID model is not suitable to capture effects of ELS on voluntary alcohol drinking.

Overall, these data indicate that ELS-altered sensitivity to the stimulatory and sedative effects of alcohol may in turn promote enhanced escalation of long-term alcohol drinking and preference.

### ELS impairs sociability, but not social novelty, following alcohol drinking in females

Because both ACEs/ELS and AUD are known to independently impair a variety of social behaviors, we reasoned that social behaviors may be a valuable readout for assessing the interactive effects of ELS and a history of EtOH drinking. To investigate these interactive effects on social behavior, CTL and ELS mice were randomly assigned to undergo either water-only or 2BC-IA alcohol drinking for three weeks. Four days after the last session of alcohol access, mice were assessed for sociability and social novelty behavior (Fig. 5A).

**Figure 5:**
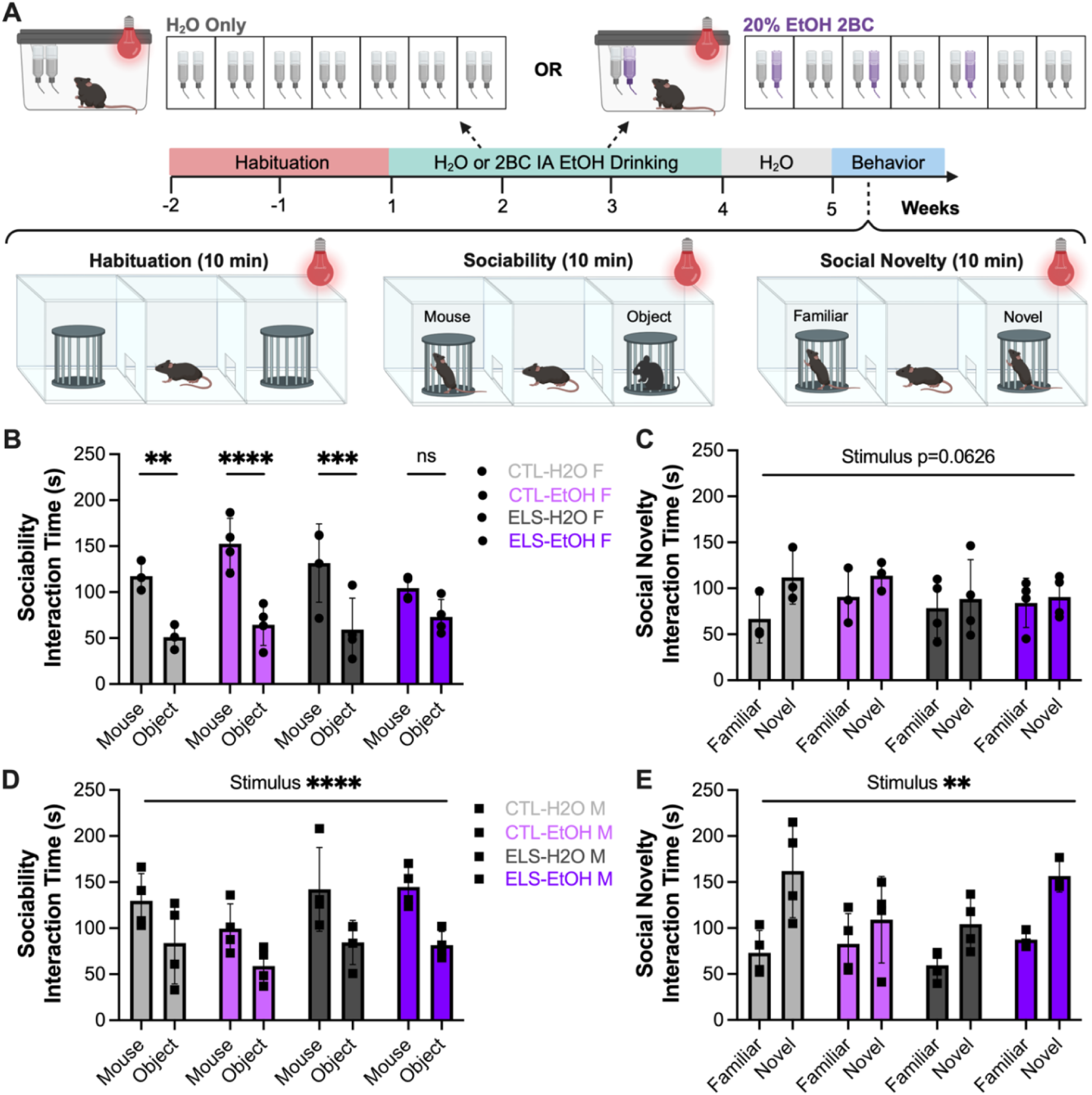
ELS impairs sociability, but not social novelty, following alcohol drinking in females. **A** Schematic of water-only or 2BC-IA alcohol drinking followed by social behavior tests. **B** Interaction time with novel mouse vs. object in CTL and ELS. Main effect of ****Stimulus (p < 0.0001, F^1, 11^ = 110.0) and *ELS x EtOH x Stimulus interaction (p = 0.0272, F^1, 11^ = 6.476). Significant difference between mouse and object interaction times in **CTL-H2O (p = 0.0020), ****CTL-ETOH (p < 0.0001), and ***ELS-H2O (p = 0.0003) females, but no difference in ELS-EtOH females. **C** Interaction time with novel vs. familiar mouse in CTL and ELS females. Trend towards a main of effect of Stimulus (p = 0.0626, F^1, 10^ = 4.390). **D** Interaction time with novel mouse vs. object in CTL and ELS. Main effect of ****Stimulus (p < 0.0001, F^1, 12^ = 53.10). Significant difference between mouse and object interaction times in *CTL-H2O (p = 0.0287), **ELS-H2O (p = 0.0063), and **ELS-ETOH (p = 0.0032) males, and a trend towards a difference in CTL-EtOH (p = 0.0571) males. **E** Interaction time with novel vs. familiar mouse in CTL and ELS males. Main effects of **Stimulus (p = 0.0012, F^1, 11^ = 18.74) and an *ELS x EtOH interaction (p = 0.0141, F^1, 11^ = 8.478). H2O: 3 CTL F, 4 ELS F, 4 CTL M, 4 ELS M EtOH: 4 CTL F, 4 ELS F, 5 CTL M, 4 ELS M

In females, both H2O- and EtOH-drinking CTL mice, as well as H2O-drinking ELS mice, interacted with the novel mouse significantly more than the novel object during the sociability test, as expected^29,30^. However, EtOH-drinking ELS females did not display a significant difference in interaction time between the mouse and object, indicating impaired sociability (Fig. 5B). Among female mice in the social novelty test, all groups tended to interact more with the novel mouse than the familiar mouse, as expected, but we did not identify any significant effects of ELS or EtOH drinking history on social novelty (Fig. 5C). In males, all groups displayed typical stimulus preferences in both the sociability and social novelty tests with no effects of ELS or EtOH drinking history (Fig. 5D-E). Overall, these results indicate that ELS and a history of EtOH drinking interact to produce sex-specific sociability deficits in females.

### Alcohol drinking may reverse the effects of ELS on anxiety-like behavior in males

To address the possibility that impaired sociability in EtOH-experienced ELS females could be primarily explained by ELS-enhanced anxiety, we tested separate groups of mice for standard anxiety-like behaviors under alcohol naïve conditions and one week after four weeks of alcohol DID. Anxiety-like behaviors were assessed in the elevated plus maze (EPM), open field test (OFT), and novelty suppressed feeding test (NSFT) as the percent of time spent in “exposed” zones (open arm, center, and food zone, respectively) (Supplemental Fig. 4A-C).

In females, ELS did not affect % open arm, % center, or % food zone time, nor locomotor activity (Supplemental Fig. 4D-F). In males, ELS mice displayed increased % open arm time relative to CTL males under alcohol naïve conditions, but low % open arm time following alcohol drinking. ELS males also displayed lower locomotor activity in the EPM than CTL males under both conditions. In the OFT and NSFT, ELS did not affect % center or % food zone, nor locomotor activity. These data suggest that ELS increases anxiety-like behavior in males with a history of alcohol drinking, specifically in the EPM (Supplemental Fig. 4G-I).

### ELS does not affect anhedonia behavior in a sucrose drinking in the dark model

Additionally, to address the possibility that impaired sociability in EtOH-experienced ELS females could be primarily explained by ELS-enhanced of depressive-like behavior, we also assessed anhedonia using a two-bottle choice limited access (2BC-LA) sucrose DID model with four 4-hour drinking sessions per week over two weeks. We did not identify any significant effects of ELS on weekly sucrose intake, sucrose preference, water intake, total liquid intake, or food intake, nor on escalation of these measures across weeks, in females or males (Supplemental Fig. 5A-J)

### ELS does not affect estrous-dependent differences in alcohol locomotor sensitivity and alcohol drinking

Lastly, to investigate whether ELS altered estrous cycle length or variability, female mice were estrous staged daily for two weeks. Females displayed an average cycle length of approximately four days, as expected^31,32^, and ELS did not affect average cycle length or range of cycle length (Supplemental Fig. 6A-C). Although ELS did not directly disrupt estrous cycling, we also sought to determine whether ELS influenced estrous-dependent changes in alcohol locomotor sensitivity and alcohol drinking. We found that females in high E2 were slightly less sensitive to the stimulatory effects of alcohol, but this trend-level effect of estrous was observed in both CTL and ELS females. Estrous stage did not affect LORR latency or duration (Supplemental Fig. 6D-G). During the first four weeks of long-term 2BC-IA alcohol drinking, we identified trends towards increased EtOH intake and preference (but not total liquid or food intake) in high E2 females, but these trend-level effects of estrous were observed in both CTL and ELS females (Supplemental Fig. 6H-K).

## DISCUSSION

### LBN+MS to investigate the effects of ELS on AUD susceptibility

Limited bedding/nesting (LBN) and maternal separation (MS) are widely used to investigate the long-term effects of early life stress (ELS) in rodents. Here, we demonstrated that the combined LBN+MS model of ELS generates fragmented maternal care, as evidenced by an increased number of nest entries/exits, and reduced body weight of pups at weaning (Fig. 1), consistent with prior reports^20,33-35^. Importantly, LBN and MS generate complementary but distinct forms of stress. LBN models resource scarcity by creating a chronically stressful environment with limited bedding and nesting materials, causing dams to provide more fragmented and unpredictable care to their pups. In contrast, MS models neglect through direct removal of the dam for extended periods of time during which pups are deprived of all maternal care (nursing, grooming, temperature regulation). Beyond differences between models, there is also significant variability within how each paradigm is implemented across laboratories. In prior studies using MS, the reported durations of separation have ranged from short 15-minute separations to full 24-hour separations, and pups may be left together during maternal separation or isolated from their littermates as well. The period of implementation has also ranged from a single day up to two weeks and has occurred at different postnatal ages within the pre-weaning period^25,26,35,36^.

Taken together, these factors have created significant variability both between and within ELS models. Combined with differences in strain, age at testing, and sex inclusion, it has been historically difficult to systematically evaluate the effects of ELS on general sex-specific behavioral outcomes, and on specific AUD-associated behaviors. Given the strong, graded association between the number of ACEs and the prevalence of self-reported alcohol problems and AUD^37-40^, we suggest that the combined LBN+MS paradigm more closely models exposure to multiple forms of childhood adversity experienced in humans with ACE-associated AUD risk.

### Behavioral sensitivity as a predictor of AUD susceptibility

Locomotor activity and loss of righting reflex are commonly used to assess the stimulatory and sedative properties, respectively, of many classes of drugs, but very few studies have examined the effects of ELS on behavioral sensitivity to alcohol in mice. We found that LBN+MS in C57BL/6J background mice increased sensitivity to the stimulatory effects of alcohol in adult females (Fig. 2), as well as decreased sensitivity to the sedative effects of alcohol in both sexes (Fig. 3). Our findings align with a prior report in which repeated MS (3 hrs/day P2-P14) increased sensitivity to the stimulatory effects of alcohol in adult female, but not male, Swiss mice^41^. Furthermore, our findings also align with a study showing that LBN (P2-P9) decreased sensitivity to the sedative effects of alcohol in female C57BL/6J mice (males not tested)^42^.

The pattern of increased stimulatory sensitivity and decreased sedation sensitivity that we observed in ELS mice closely matches the “modified differentiator model of alcohol response” phenotypes that are predictive of AUD risk in humans^43^. Longitudinal studies performed in both men and women by the Chicago Social Drinking Project reported increased alcohol stimulation and decreased alcohol sedation experienced by heavy drinkers relative to light drinkers^43,45^, as well as a persistent inverse association between stimulatory and sedative effects (i.e. people who experienced stronger stimulant effects were more likely to experience lower sedative effects)^46^. Prior work also demonstrated that higher sensitivity to alcohol stimulation and lower sensitivity to alcohol sedation robustly predicts increased AUD symptoms, including increased binge drinking, up to 10 years later^14,45,47^.

### Effects of ELS on alcohol drinking

We next investigated the effects of LBN+MS on voluntary alcohol drinking. Using a long-term intermittent access two-bottle choice drinking model, we found that ELS promotes escalation of alcohol drinking and preference, particularly in males (Fig. 4). Very few studies have examined the effects of ELS on voluntary alcohol drinking in mice, and the few existing studies examined drinking behaviors only in adolescent males^48^ or in combination with adolescent social isolation stress^49^ or forced alcohol dependence via chronic intermittent ethanol (CIE) vapor inhalation^34^. In one study using short-term 2BC limited access 15% EtOH DID (2 hrs/day x 2 weeks), LBN (P2-P9) accelerated escalation of alcohol intake in males, but not females^34^, similar to the pattern we identified in our long-term 2BC-IA paradigm. We chose not to perform within-subject testing of sequential alcohol motor sensitivity and alcohol drinking in the same mice, due to the potential for carryover effects.

### Effects of ELS on social and affective behaviors

In addition to escalated alcohol intake, AUD is associated with a variety of other behavioral changes, including social behavior, anxiety, and depression. We found that a combination of LBN+MS and a history of alcohol drinking (three weeks of 2BC-IA followed by four days of abstinence) produced deficits in sociability, but not social novelty, only in females (Fig. 5). A previous study using 2BC limited access alcohol drinking (three weeks of DID followed by seven days of abstinence) in non-ELS mice identified a female-specific effect in which a history of alcohol drinking decreased social novelty preference^30^. These findings suggest that distinct drinking schedules or withdrawal periods may differentially exacerbate social deficits, and that female mice may be particularly susceptible to the adverse effects of alcohol and/or ELS on social behaviors. However, MS (P2-P5) and early weaning decreased social novelty in adolescent male mice (females not tested)^48^, suggesting that males may also be sensitive to ELS-induced social deficits at specific developmental timepoints (i.e., adolescence). Additionally, while there is an abundance of literature examining the effects of ELS on affective behaviors, there are no clear, consistent phenotypes of anxiety-like behavior or anhedonia with or without prior alcohol exposure^25,50,51^.

### Extended amygdala CRF circuitry as a possible mediator of ELS effects on AUD susceptibility

Substantial clinical evidence indicates that ACEs disrupt normal development of stress-sensitive corticotropin-releasing factor (CRF) circuitry in humans, resulting in elevated cerebrospinal fluid CRF levels and dysregulated blood cortisol levels in adulthood^52^. In C57BL/6 mice, ELS upregulated CRF expression in stress and reward circuitry regions^33,53-55^, and inhibition of CRF neurons within the central and basolateral amygdala rescue ELS-induced effects on threat reactivity and reward seeking behaviors^56,57^. The CRF system within the extended amygdala network is also implicated in the response to stress within the context of excessive alcohol consumption^58,59^, suggesting that ELS may induce lasting impacts on alcohol sensitivity and drinking via plasticity in these circuits. Like other forebrain structures, maturation of the bed nucleus of the stria terminalis (BNST), a sexually dimorphic region in the extended amygdala^60-64^, is incomplete during early postnatal development^65^, making it particularly susceptible to disruption by ELS. Additionally, dorsolateral nuclei of the BNST are a prominent source of extra-hypothalamic CRF. Given that activation of BNST-CRF neurons is implicated in other AUD-associated behaviors, particularly binge alcohol drinking^66-69^, this circuitry is a promising potential mediator of ELS-modulated alcohol sensitivity and AUD susceptibility.

### Conclusions

In this study, we implemented and validated a robust LBN+MS model to investigate the effects of ELS on AUD-associated behaviors in adult female and male mice. We found that ELS increased sensitivity to the stimulatory effects of alcohol, particularly in females, and decreased sensitivity to the sedative effects of alcohol, particularly in males, closely mirroring the “modified differentiator model” of AUD susceptibility observed in humans^43^. We also found that ELS increased escalation of voluntary alcohol drinking in males and disrupted sociability specifically in females with a history of alcohol drinking. Future work should investigate the neurobiological mechanisms mediating these observed effects of ELS on alcohol behavioral sensitivity and drinking, and we propose BNST-CRF circuitry as a promising candidate. Additionally, given the associations between the number of ACEs and prevalence of other substance usage (psychostimulants, opioids, etc.)^40,70^, we hypothesize that assessments of behavioral locomotor sensitivity may be a promising generalizable strategy for first-pass investigations ELS impacts on susceptibility to other substance use disorders beyond AUD.

## ACKNOWLEDGEMENTS

This work was supported by funding to A.R.M (NIH/NIAAA F31 AA031630), O.S.R.L. (Stanford Neuroscience Undergraduate Research Opportunity), A.C.Y. (Stanford Maternal Child Health Research Institute), and W.J.G. (NIH/NIAAA R00 AA025677, Stanford WHSDM, Stanford NeuroChoice, Stanford Taube Youth Addiction Initiative, Whitehall Foundation, Brain Research Foundation). We sincerely thank Dr. Yihe Ma and Emmalyn Leonard for contributions to initial experiments. We thank Dr. Belgin Yalcin, Jerome Geronimo, and Stanford Veterinary Service Center staff for technical assistance. We extend our gratitude to all Giardino Lab members and Drs. Karen Parker, Scott Owen, Andrea Goldstein-Piekarski, Xiaoke Chen, Matthew Pomrenze, Joseph Garner, and Kevin Bath for invaluable feedback and discussion.

## SUPPLEMENTAL FIGURES

**Supplemental Figure 1:**
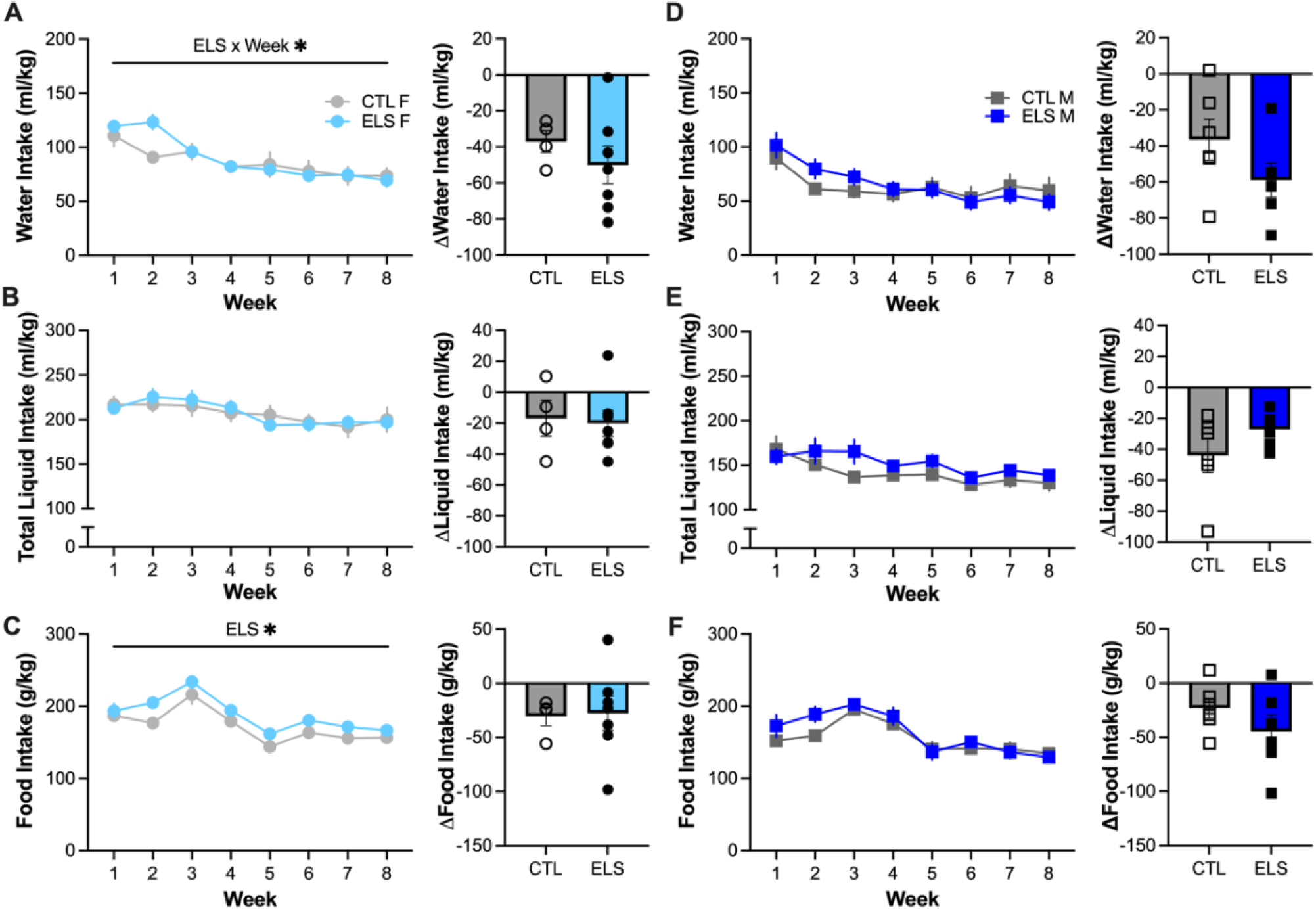
ELS does not affect escalation of water, total liquid, or food intake in the long-term 2BC-IA drinking model. **A** Left: Water intake in CTL and ELS females. Main effects of ****Week (p < 0.0001, F^3.933, 37.08^ = 15.22) and an *ELS x Week interaction (p = 0.0414, F^7, 66^ = 2.243). Significant difference between CTL and ELS in *Week 2 (p = 0.0299). Right: Difference in water intake between Week 8 and Week 1. No difference between groups. **B** Left: Total liquid intake in CTL and ELS females. Main effect of **Week (p = 0.0022, F^3.233, 30.48^ = 5.920). Right: Difference in total liquid intake between Week 8 and Week 1. No difference between groups. **C** Left: Food intake in CTL and ELS females. Main effects of *ELS (p = 0.0160, F^1, 10^ = 8.378) and ****Week (p < 0.0001, F^3.399, 31.56^ = 24.49). Right: Difference in food intake between Week 8 and Week 1. No difference between groups. **D** Left: Water intake in CTL and ELS males. Main effect of ****Week (p < 0.0001, F^2.775, 35.68^ = 17.43). Right: Difference in water intake between Week 8 and Week 1. No difference between groups. **E** Left: Total liquid intake in CTL and ELS males. Main effect of ****Week (p < 0.0001, F^3.374, 43.38^ = 10.58). Right: Difference in total liquid intake between Week 8 and Week 1. No difference between groups. **F** Left: Food intake in CTL and ELS males. Main effect of ****Week (p < 0.0001, F^3.638, 46.25^ = 33.81). Right: Difference in food intake between Week 8 and Week 1. No difference between groups.

**Supplemental Figure 2:**
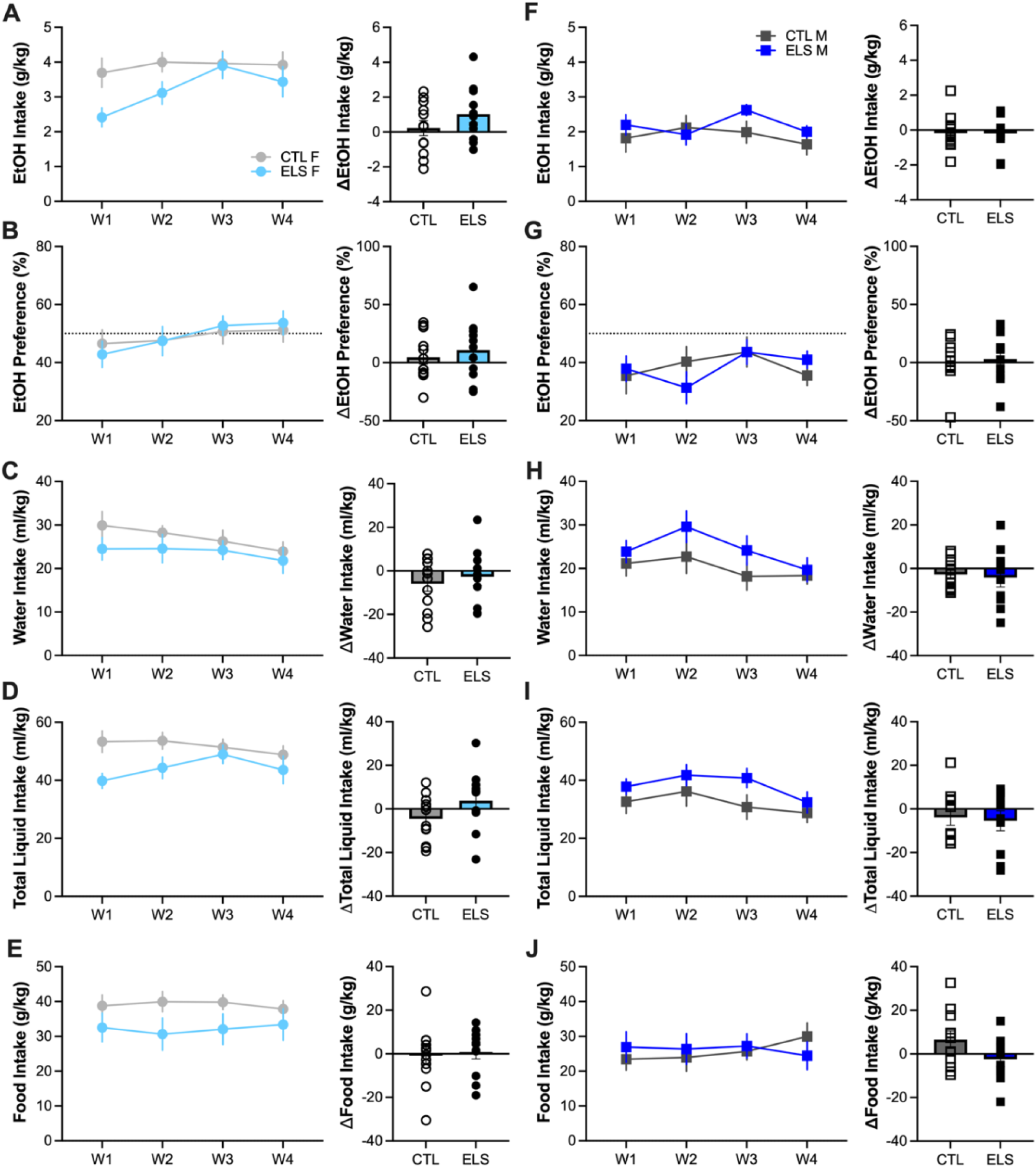
ELS does not affect alcohol intake or preference in the short-term 2BC-LA DID model. **A** Left: Ethanol intake in CTL and ELS females. Main effect of *Week (p = 0.0164, F^2.273, 47.74^ = 4.240). Right: Difference in ethanol intake between Week 4 and Week 1. No difference between groups. **B** Left: Ethanol preference in CTL and ELS females. No main effect of ELS or Week. Right: Difference in ethanol preference between Week 4 and Week 1. No difference between groups. **C** Left: Water intake in CTL and ELS females. No main effect of ELS or Week. Right: Difference in water intake between Week 4 and Week 1. No difference between groups. **D** Left: Total liquid intake in CTL and ELS females. No main effect of ELS or Week. Right: Difference in total liquid intake between Week 4 and Week 1. No difference between groups. **E** Left: Food intake in CTL and ELS females. No main effect of ELS or Week. Right: Difference in food intake between Week 4 and Week 1. No difference between groups. **F** Left: Ethanol intake in CTL and ELS males. No main effect of ELS or Week. Right: Difference in ethanol intake between Week 4 and Week 1. No difference between groups. **G** Left: Ethanol preference in CTL and ELS males. No main effect of ELS or Week. Right: Difference in ethanol preference between Week 4 and Week 1. No difference between groups. **H** Left: Water intake in CTL and ELS males. Main effect of *Week (p = 0.0183, F^2.210, 42^ = 4.222). Right: Difference in water intake between Week 4 and Week 1. No difference between groups. **I** Left: Total liquid intake in CTL and ELS males. Main effect of *Week (p = 0.0101, F^2.536, 48.18^ = 4.532). Right: Difference in total liquid intake between Week 4 and Week 1. No difference between groups. **J** Left: Food intake in CTL and ELS males. No main effect of ELS or Week Right: Difference in food intake between Week 4 and Week 1. No difference between groups. n = 12 CTL F, 11 ELS F, 12 CTL M, 10 ELS M

**Supplemental Figure 3:**
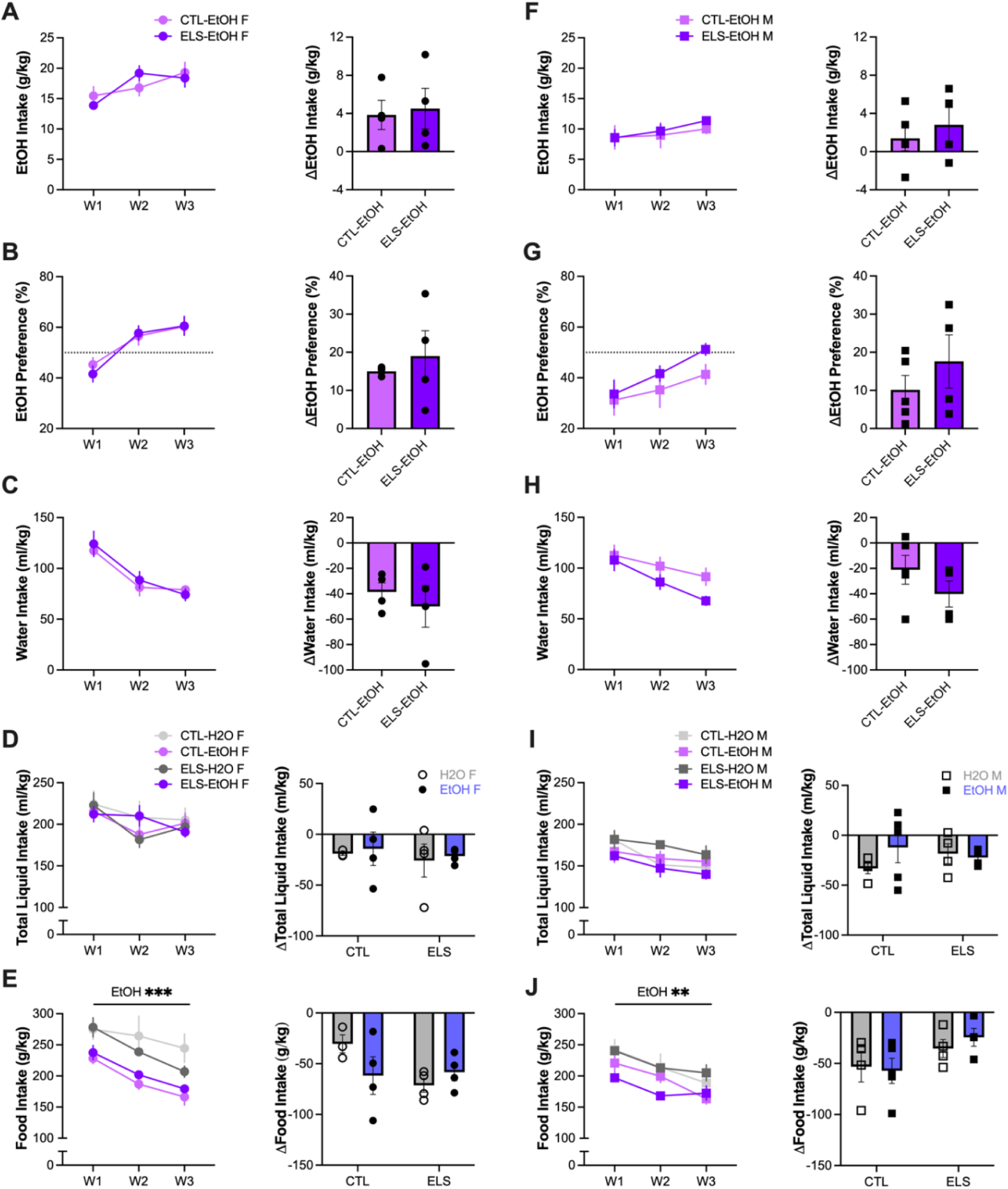
ELS does not affect alcohol intake or preference in only three weeks of 2BC-IA drinking. **A** Left: Ethanol intake in CTL and ELS females that underwent 2BC-IA ethanol drinking. Main effect of *Week (p = 0.0337, F^1.766, 10.59^ = 4.936). Right: Difference in ethanol intake between Week 3 and Week 1. No difference between groups. **B** Left: Ethanol preference in CTL and ELS females that underwent 2BC-IA ethanol drinking. Main effect of **Week (p = 0.0011, F^1.753, 10.52^ = 14.78). Right: Difference in ethanol preference between Week 3 and Week 1. No difference between groups. **C** Left: Water intake in CTL and ELS females that underwent 2BC-IA ethanol drinking. Main effect of **Week (p = 0.0074, F^1.075, 6.451^ = 14.37). Right: Difference in water intake between Week 3 and Week 1. No difference between groups. **D** Left: Total liquid in CTL and ELS females that underwent 2BC-IA ethanol drinking or water-only drinking. Main effect of *Week (p = 0.0113, F^1.708, 18.79^ = 6.159). Right: Difference in total liquid intake between Week 3 and Week 1. No main effect of ELS or EtOH. **E** Left: Food intake in CTL and ELS females that underwent 2BC-IA ethanol drinking or water-only drinking. Main effects of ***EtOH (p = 0.0007, F^1, 11^ = 21.53) and ****Week (p < 0.0001, F^1.407, 15.48^ = 45.26). Right: Difference in food intake between Week 3 and Week 1. No main effect of ELS or EtOH. **F** Left: Ethanol intake in CTL and ELS males that underwent 2BC-IA ethanol drinking. No main effect of ELS or Week. Right: Difference in ethanol intake between Week 3 and Week 1. No difference between groups. **G** Left: Ethanol preference in CTL and ELS males that underwent 2BC-IA ethanol drinking. Main effect of **Week (p = 0.0040, F^1.686, 11.80^ = 9.899). Right: Difference in ethanol preference between Week 3 and Week 1. No difference between groups. **H** Left: Water intake in CTL and ELS males that underwent 2BC-IA ethanol drinking. Main effect of **Week (p = 0.0028, F^1.745, 12.22^ = 10.48). Right: Difference in water intake between Week 3 and Week 1. No difference between groups. **I** Left: Total liquid in CTL and ELS males that underwent 2BC-IA ethanol drinking or water-only drinking. Main effect of ***Week (p = 0.0008, F^1.931, 25.11^ = 9.786). Right: Difference in total liquid intake between Week 3 and Week 1. No main effect of ELS or EtOH. **J** Left: Food intake in CTL and ELS males that underwent 2BC-IA ethanol drinking or water-only drinking. Main effects of **EtOH (p = 0.0089, F^1, 13^ = 9.43) and ****Week (p < 0.0001, F^1.975, 25.67^ = 23.63). Right: Difference in food intake between Week 3 and Week 1. Trend towards a main effect of ELS (p = 0.0514, F^1, 13^ = 4.601).

**Supplemental Figure 4:**
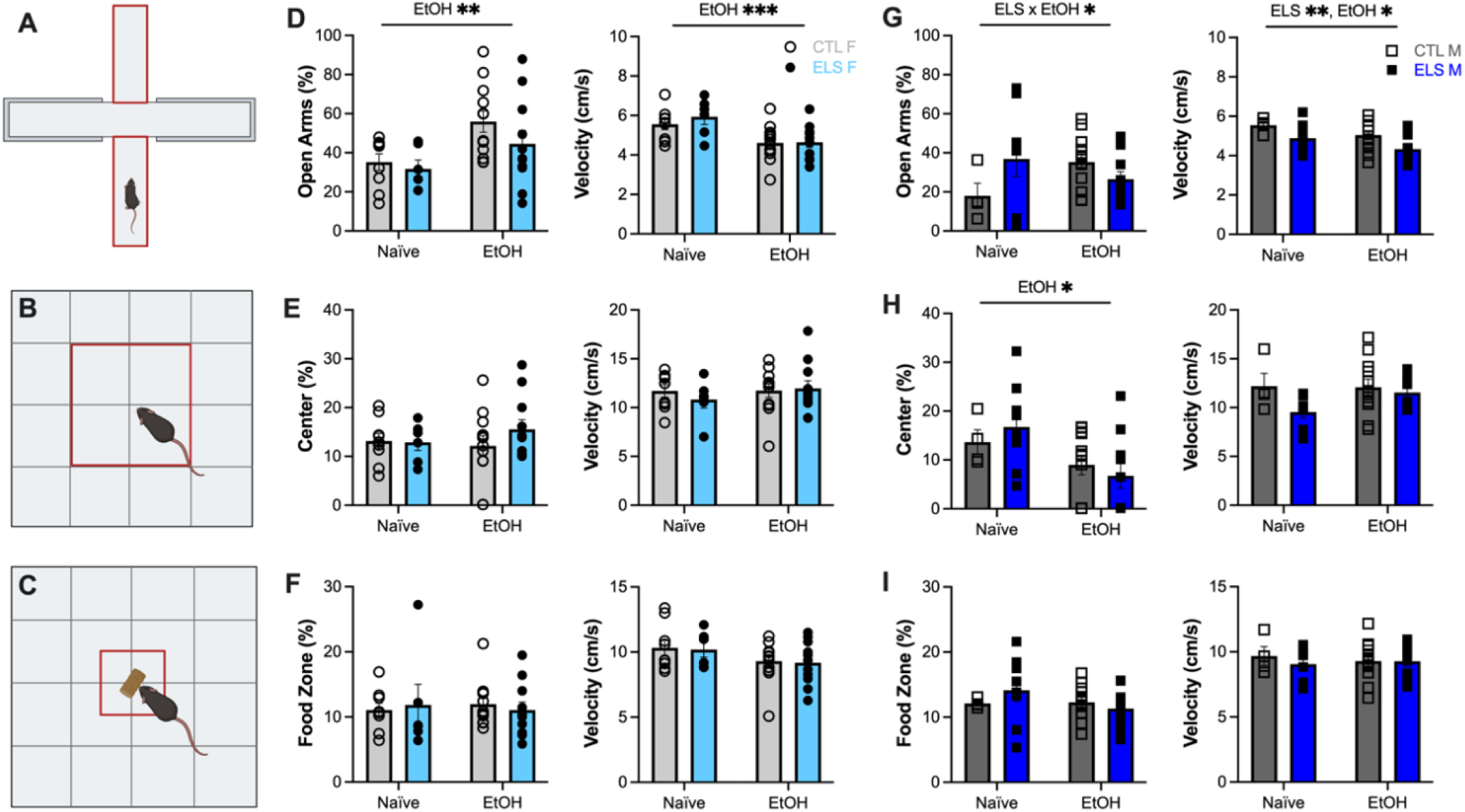
Alcohol drinking may reverse the effect of ELS on anxiety-like behavior in males, particularly in the elevated plus maze. **A** Elevated Plus Maze with Open Arms outlined in red. **B** Open Field Test with Center outlined in red. **C** Novelty Suppressed Feeding Test with Food Zone outlined in red. **D** Left: Time spent in the Open Arms as a percentage of total trial time in CTL and ELS females. Main effect of **EtOH (p = 0.0091, F^1, 34^ = 7.663). Right: Average velocity across the trial. Main effect of ***EtOH (p = 0.0006, F^1, 34^ = 14.35). **E** Left: Time spent in the Center as a percentage of total trial time in CTL and ELS females. No main effect of ELS or EtOH. Right: Average velocity across the trial. No main effect of ELS or EtOH. **F** Left: Time spent in the Food Zone as a percentage of total trial time in CTL and ELS females. No main effect of ELS or EtOH. Right: Average velocity across the trial. No main effect of ELS or EtOH. **G** Left: Time spent in the Open Arms as a percentage of total trial time in CTL and ELS males. Main effect of *ELS x EtOH interaction (p = 0.0423, F^1, 31^ = 4.487). Right: Average velocity across the trial. Main effects of **ELS (p = 0.0095, F^1, 31^ = 7.642) and *EtOH (p = 0.0413, F^1, 31^ = 4.532). **H** Left: Time spent in the Center as a percentage of total trial time in CTL and ELS males. Main effect of *EtOH (p = 0.0138, F^1, 31^ = 6.822). Right: Average velocity across the trial. No main effect of ELS or EtOH. **I** Left: Time spent in the Food Zone as a percentage of total trial time in CTL and ELS males. No main effect of ELS or EtOH. Right: Average velocity across the trial. No main effect of ELS or EtOH. Naïve: n = 9 CTL F, 6 ELS F, 4 CTL M, 9 ELS M EtOH: n = 12 CTL F, 11 ELS F, 12 CTL M, 10 ELS M

**Supplemental Figure 5:**
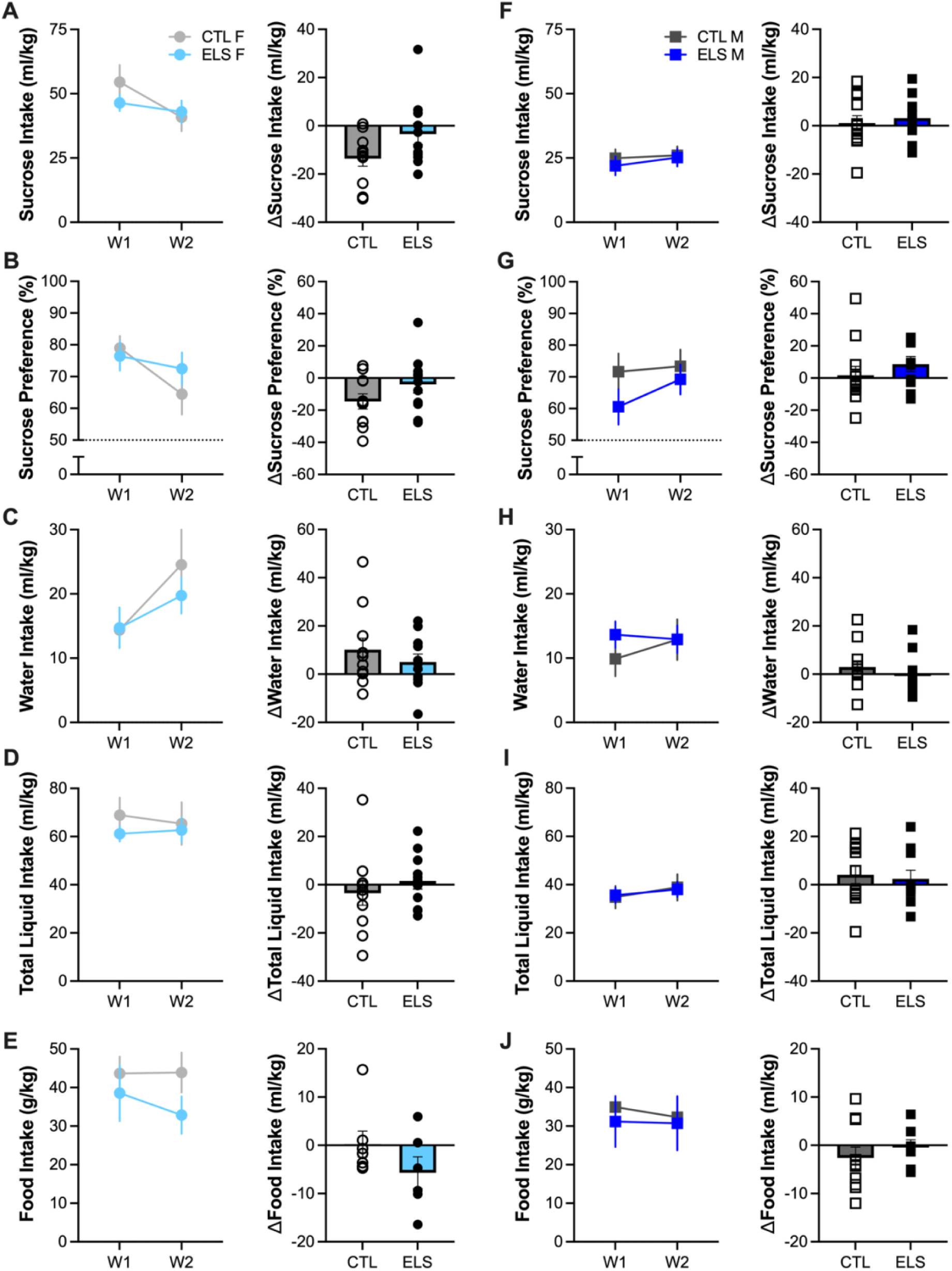
ELS does not affect sucrose intake or preference in the short-term 2BC-LA DID model. **A** Left: Sucrose intake in CTL and ELS females. Main effect of **Week (p = 0.0045, F^1, 20^ = 10.24). Right: Difference in sucrose intake between Week 2 and Week 1. No difference between groups. **B** Left: Sucrose preference in CTL and ELS females. Main effect of *Week (p = 0.0171, F^1, 20^ = 6.768). Right: Difference in sucrose preference between Week 2 and Week 1. No difference between groups. **C** Left: Water intake in CTL and ELS females. Main effect of *Week (p = 0.0171, F^1, 20^ = 6.77). Right: Difference in water intake between Week 2 and Week 1. No difference between groups. **D** Left: Total liquid intake in CTL and ELS females. No main effect of ELS or Week. Right: Difference in total liquid intake between Week 2 and Week 1. No difference between groups. **E** Left: Food intake in CTL and ELS females. No main effect of ELS or Week. Right: Difference in food intake between Week 2 and Week 1. No difference between groups. **F** Left: Sucrose intake in CTL and ELS males. No main effect of ELS or Week. Right: Difference in sucrose intake between Week 2 and Week 1. No difference between groups. **G** Left: Sucrose preference in CTL and ELS males. No main effect of ELS or Week (. Right: Difference in sucrose preference between Week 2 and Week 1. No difference between groups. **H** Left: Water intake in CTL and ELS males. No main effect of ELS or Week. Right: Difference in water intake between Week 2 and Week 1. No difference between groups. **I** Left: Total liquid intake in CTL and ELS males. No main effect of ELS or Week. Right: Difference in total liquid intake between Week 2 and Week 1. No difference between groups. **J** Left: Food intake in CTL and ELS males. No main effect of ELS or Week. Right: Difference in food intake between Week 2 and Week 1. No difference between groups. n = 11 CTL F, 11 ELS F, 12 CTL M, 10 ELS M

**Supplemental Figure 6:**
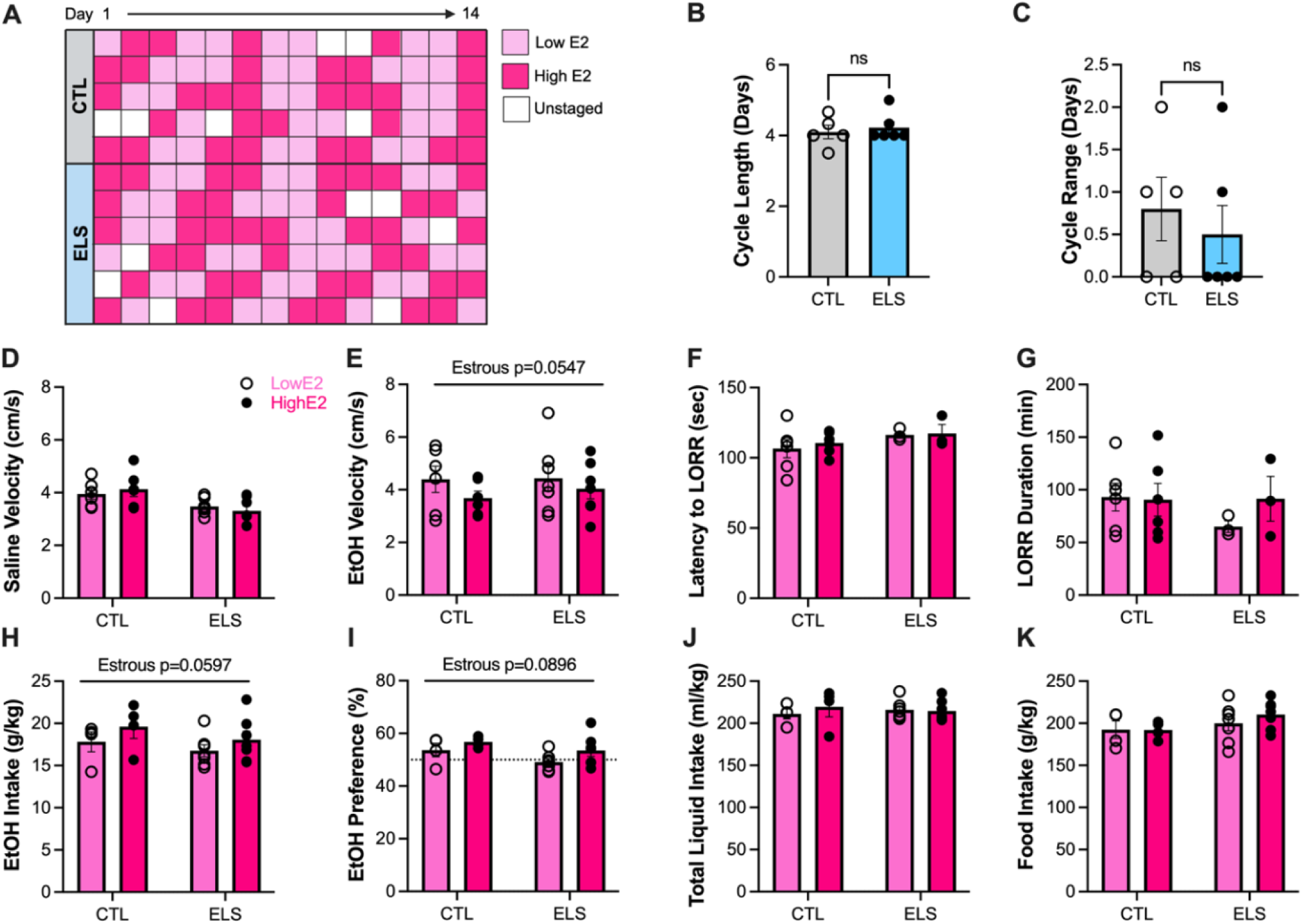
ELS does not affect estrous-dependent differences in behavioral sensitivity to alcohol and alcohol drinking. **A** Heatmap of daily estrous staging over two weeks. Each row represents one mouse and each column represents one day. **B** Average estrous cycle length. No difference between groups. **C** Range of estrous cycle length. No difference between groups. **D** Average velocity following saline i.p injection in CTL and ELS females in low or high E2. Main effect of *ELS (p = 0.0242, F^1, 11^ = 6.819). **E** Average velocity following stimulatory dose EtOH i.p injection in CTL and ELS females in low or high E2. Trend towards a main effect of Estrous (p = 0.0547, F^1, 11^ = 4.621). **F** Average latency to lose righting reflex following sedative dose EtOH i.p. injection in CTL and ELS females in low or high E2. No main effect of Estrous or ELS. **G** Average duration of loss of righting reflex in CTL and ELS females in low or high E2. No main effect of Estrous or ELS. **H** Average ethanol intake across 4 weeks of 2BC-IA alcohol drinking in CTL and ELS females in low or high E2. Trend towards a main effect of Estrous (p = 0.0597, F^1, 9^ = 4.636). **I** Average ethanol preference across 4 weeks of 2BC-IA alcohol drinking in CTL and ELS females in low or high E2. Trend towards a main effect of Estrous (p = 0.0896, F^1, 9^ = 3.618). **J** Average total liquid intake across 4 weeks of 2BC-IA alcohol drinking in CTL and ELS females in low or high E2. No main effect of Estrous or ELS. **K** Average food intake across 4 weeks of 2BC-IA alcohol drinking in CTL and ELS females in low or high E2. No main effect of Estrous or ELS. A-C: n = 5 CTL F, 6 ELS F D-E: n = 6 CTL F, 7 ELS F F-G: n = 6 CTL F, 3 ELF H-K: n = 4 CTL F, 7 ELS F

